# A crucial role for PIH1D1 in modulating p53 stability via the R2TP complex

**DOI:** 10.64898/2026.02.07.704515

**Authors:** Dhiraj Kumar Singh, Qulsum Akhter Mir, Riyaz Ahmad Mir

## Abstract

The R2TP is a multimeric protein complex consists of RUVBL1, RUVBL2, PIH1D1, and RPAP3, and it is known to functions as a specialized co-chaperone. We hypothesize that PIH1D1 recognizes p53 and stabilizes it via R2TP complex. Upon successful completion of this study, innovative mechanism has been found for the interaction and stabilization of p53 and hence govern the cell cycle. Upon interaction between p53 and PIH1D1 protein, p53 is stabilized by PIH1D1 protein, without affecting its C-terminal domain. We have also observed that p53 protein levels were affected after the alteration in expression levels of PIH1D1. Based on the finding, we suggest that the R2TP complex stabilizes and regulates P53. Therefore, this novel method will work as a flashpoint to restore the function of p53 in cancer cells, controlling cancer and cell cycle progression.

## Introduction

The protein, commonly referred to as “P53” and recognized as the guardian of the genome [1], serves to detect DNA damage, oncogene activation, telomere erosion, hypoxia, heat shock, or ribosomal stress; p53 undergoes a process of stabilization and activation[2,3]. Despite the fundamental importance of p53 in cancer biology, the precise mechanisms governing its stability, especially within the context of cancer, remain the focus of rigorous investigation. One of the important questions that remains complex is how p53 function is regulated? The R2TP complex, also called PAQosome, is the first multi-molecular co-chaperone of HSP90[4,5]. The R2TP complex is characterized by a distinct molecular architecture that reflects its intricate role as a cochaperone in various cellular processes[6]. At its core, the complex is composed of four essential subunits: RU-VBL1, RUVBL2, RPAP3, and PIH1D1. RUVBL1 and RUVBL2 are ATPases belonging to the AAA+ superfamily, providing the energy necessary for the complex’s chaperone function[7]. RPAP3 serves as a bridging component that connects the RUVBL proteins with PIH1D1. PIH1D1, in turn, acts as a scaffold, facilitating the interaction between the R2TP complex and its client proteins. Clients’ proteins are brought to R2TP through bridging proteins[8]. These bridging proteins bind to the PIH1D1 component of the R2TP complex through a motif DSDD/E, which is phosphorylated by CK2[9,10].

The R2TP complex plays a pivotal role in orchestrating the DNA damage response, influencing the stability of all PIKKs[11]. Notably, the mutation of the phospho-binding domain PIH-N resulted in diminished p53 activation post-induction of DNA double-strand breaks, even in the presence of normal levels of ATM and DNA-PKcs, two key PIKKs crucial for responding to this form of DNA damage [12]. It has been proposed that among various bridging proteins, ECD may act as a bridge[13,14]. There is currently no study that describes the interaction between the R2TP complex and P53. In this study, we sought the relationship between the R2TP complex and P53. Notably, our specific interest in exploring the interaction between the R2TP complex and p53 has yielded significant insights. We observed a direct interaction between the PIH1D1 component of the R2TP complex and p53, regardless of p53 phosphorylation status. To explore into the functional implications of this interaction, we utilized siRNA-mediated silencing of PIH1D1 expression. Intriguingly, the depletion of PIH1D1 resulted in a notable reduction in endogenous wild-type p53 levels, underscoring the crucial role of PIH1D1 in maintaining p53 stability.

## Material and Methods

### Cloning and Expression of PIH1D1 as a GST Fusion Protein

The pGEX2T vector was used for the cloning of our gene of interest PIH1D1. The vector is resistant to ampicillin and is inducible with IPTG. Construct for GST-fused PIH1D1 was generated using pGEX2T (AddGene) plasmid, and amplification of the target gene was done by the PCR technique. Restriction sites were introduced with primer sequences, which were used for the PCR. Amplified PCR product was digested with restriction enzymes, and eluted. The digested PCR product was ligated into the pGEX2T vector and transformed into an E. coli competent cell DH5*α* alpha. Several of the transformants were grown in a culture plate at 37°C. To check for expression of the target protein, we picked a colony from the culture plate. We grew the selected colony in LB media containing ampicillin, purified the plasmid, and then transformed it into BL21 competent cells. Positive expression was confirmed using induction of BL21 transformants with IPTG, and Bacterial cells were lysed to check protein expression with SDS protein gel.

### Purification and Quantification of GST-tagged PIH1D1

Resuspend the pelleted E. coli cells in 15 mL cold lysis buffer and lyse the cells by sonica-tion on ice (∼10 times for 10 sec each with 1 min rest between bursts to minimize sample heating). The lysate was centrifuged at 48,000 × g for 20 min at 4°C. Decant the supernatant into a clean 50 ml centrifuge tube. Pellet was resuspended in a 15-ml PBS buffer using a Dounce homogenizer. 5–10 *µ*l each of lysate, supernatant, and resuspended pellet were analysed by SDS-PAGE gel to verify that the fusion protein is in the supernatant fraction. The soluble fusion protein was loaded onto the equilibrated glutathione Sepharose column using a flow rate of 0.1 ml/min. Fractions were collected and run on gels to verify that the fusion protein is binding to the column and that the capacity has not been exceeded Pool fractions that contain the GST fusion protein.

### In Vitro binding Assay: GST-tagged PIH1D1

To determine the region of p53 interaction with PIH1D1, the full length of p53, CTD-lacking p53 (w/o CTD), and the CTD-only p53 fragment were cloned into the PGEX2T plasmid having a GST tag. Proteins were expressed in vitro and purified with glutathione-S-transferase (GST) beads (Fig. 4A). Quantification of GST-tagged P53 protein fragments was done with SDS-PAGE, and their (CTD, W/O CTD p53, and full-length p53) interaction with PIH1D1 was performed with the help of MCF7 cell lysates. The results were analysed using Western blots.

### Cell Culture and Transfection

MCF7 and HeLa (NCCS, Pune) was grown in in Dulbecco’s Modified Eagle Medium (HiMedia, Cat. AL007A) and supplemented with 10% Fetal Bovine Serum and Serum (Gibco, Cat. 10270106) and Antibiotic-Antimycotic100X (Gibco, Cat. 15240096). All cells were grown in an atmosphere of 5% CO2 at 37°C and were sub-cultured by trypsinization with Trypsin - EDTA Solution 1X (HiMedia, Cat. TCL014). Cells were harvested and lysed in urea lysis buffer [7M urea, 20mM HEPES (pH 7.6), 25mM NaCl, 0.05% (v/v) Triton X-100, 0.1M dithiothreitol, 5mM NaF, 2mM Na□VO□, 2.5mM Na□P□O□, and 1 x Complete Mini Protease Inhibitor Cocktail (Roche Diagnostics, Burgess Hill, UK)]. The cells were incubated on ice for 30 minutes, followed by centrifugation at 13,000 g for 10 minutes at 4°C.

A verified shRNA construct (Catalogue No. TL302919, OriGene Technologies, USA) was used to knock down PIH1D1 (Gene ID: 55011) in cells cultivated under normal conditions by RNA interference. A negative control was a non-targeting shRNA. Using a commercially available reagent and the manufacturer’s procedure tailored for the cell line, transfection was performed. Antibiotic selection was used where appropriate to find populations that expressed shRNA. For PIH1D1 transcripts, the effectiveness of the knockdown was assessed by quantitative RT-PCR, and immunoblotting was used to confirm the protein level. Several shRNA sequences from the catalog were examined to guarantee specificity and reduce off-target effects, and rescue studies using a shRNA-resistant PIH1D1 construct were used to confirm phenotypes. For the knockdown and control groups, functional tests such as cell viability, cell-cycle analysis, and evaluation of R2TP complex-related signalling were carried out concurrently.

### Western blotting

Equal amounts of protein (20□µg) were loaded on a 4–20% polyacrylamide gel (miniprotean TGX precast protein gels; Bio-rad) and electrophoretically separated in running buffer [25□mM Tris, 192□mM glycine, 0.1% SDS, H2O.] at a constant current of 100□V (Bio-rad Mini-PROTEAN® Tetra System). After electrophoresis with SDS-PAGE, the seprated proteins were transferred onto a PVDF membrane (0.45-μm pore size; Merck Millipore) at 300 mA for 60 min in ice-cold 20% (vol/vol) methanol transfer buffer. After transfer, the membrane was stained with Ponceau S (SRL, Cat.38610) and washed with distilled water using a shaker The membrane was blocked with 5% non-fat milk powder (HiMedia, Cat. GRM1254) or 3% Bovine Serum Albumin/BSA (SRL, Cat.85171) in 1X TBS-T buffer for 1 hour at room temperature. The membrane was incubated overnight with the primary antibodies. Then membranes were therefore exposed to HRP-conjugated anti-mouse secondary antibody or anti-rabbit secondary antibody 1□h at room temperature. Proteins were visualized using Thermo Scientific TM Pierce TM ECL Western Blotting Substrate (Cat. 32109) or G-Biosciences femtoLUCENT TM PLUS-HRP (Cat. 786-081) and analysed by Azure Imaging Systems (Azure Biosystems, Sierra Ct, Dublin, CA, USA). Anti-GAPDH was used as a housekeeping protein to normalise the integrated intensities.

### Cell cycle analysis

Briefly, 3 × 10^5^ of MCF-7 cells were seeded in each well of a 6-well plate and treated with EADs at 25 and 50 μg/mL. Control untreated cells were also included. After incubation for 24, 48, and 72 hours, the cells were trypsinized and washed with PBS. After centrifugation, the cell suspension was resuspended repeatedly in single cells prior to fixation with 70% ethanol. Fixed cells were kept at -20°C for at least 2 hours. Later, fixed cells were washed with PBS twice, and the supernatant was discarded. Cell pellets were resuspended with 425 μL of PBS in a round-bottom tube. Next, 50 μL of RNAse and 25 μL of propidium iodide were added to the cell suspension and incubated for 15 minutes on ice in the dark. FACS Calibur (BD Biosciences, USA) and Cell Quest Pro software (BD Biosciences, USA) were used to determine the cell cycle distribution. A total of 10,000 cells were acquired each time using a FACS Calibur flow cytometer. Flow cytometric data were analysed using Modfit software and displayed in a histogram of cell count (y-axis) against DNA content (*x*-axis).

### Quantitative real-time PCR

For quantitative analysis of gene expression levels, RNA was isolated using TRI Reagent solution (sigma) and 2 μg of total RNA was reverse transcribed using random hexamers. An aliquot containing 200 ng of the total cDNA was subjected to real-time PCR using primers specific for p21 and p53. Primers were designed using the Primer-BLAST software. Real-time PCR reactions were carried out in a 20-μL total volume containing 1X SYBR Green qPCR Master Mix (Thermo Scientific, Waltham, USA). PCR conditions comprised 40 cycles of denaturation at 94°C for 30 s, annealing at 59°C for 45 s, extension at 72°C for 1 min. Similarly, cDNA was also amplified using primers specific for GAPDH, which served as the internal control. The amplification of the expected sizes of PCR products was confirmed by running them on an agarose gel (1.5%). Cycle threshold (Ct) values were calculated for each PCR, and relative quantification was calculated using 2–ΔΔCt method.

### Statistical analysis

While quantitative real-time PCR findings were evaluated using the ΔΔCq method with housekeeping genes as internal controls, Western blot data were quantified by densitometry and normalized to the relevant loading control. Relative expression was determined as 2^(-ΔΔCq). Technical duplicates were averaged prior to statistical analysis, which used data from independent biological replicates (n indicated in figure legends). An unpaired two-tailed Student’s t-test was used for comparisons between two groups, and a one-way ANOVA was used for comparisons between several groups. When feasible, individual data points are displayed alongside the mean ± SEM, and p < 0.05 was deemed statistically significant. GraphPad Prism was used for all analysis and graph production.

## Results

### PIH1D1 Acts as a Link Between p53 and the R2TP Complex

In order to answer this question whether p53 is directly interact with P1H1D1 or does it need a bridging protein component of R2TP complex. For this purpose, the expression and purification of recombinant GST-tagged PIH1D1 protein were confirmed through SDS–PAGE, Coomassie staining, and Western blot analyses. GST-PIH1D1 pulldown was carried out using pure recombinant p53 protein alone or the pure protein was incubated with HeLa cell lysate (Note: HeLa cell does not have p53 protein present under normal conditions due to HPV E6). Notably GST-PIH1D1 was able to pull down pure p53 both alone as well as from the cell lysate of Hela cells in which purified p53 was incubated (Figure 1A).

**Figure 1.**
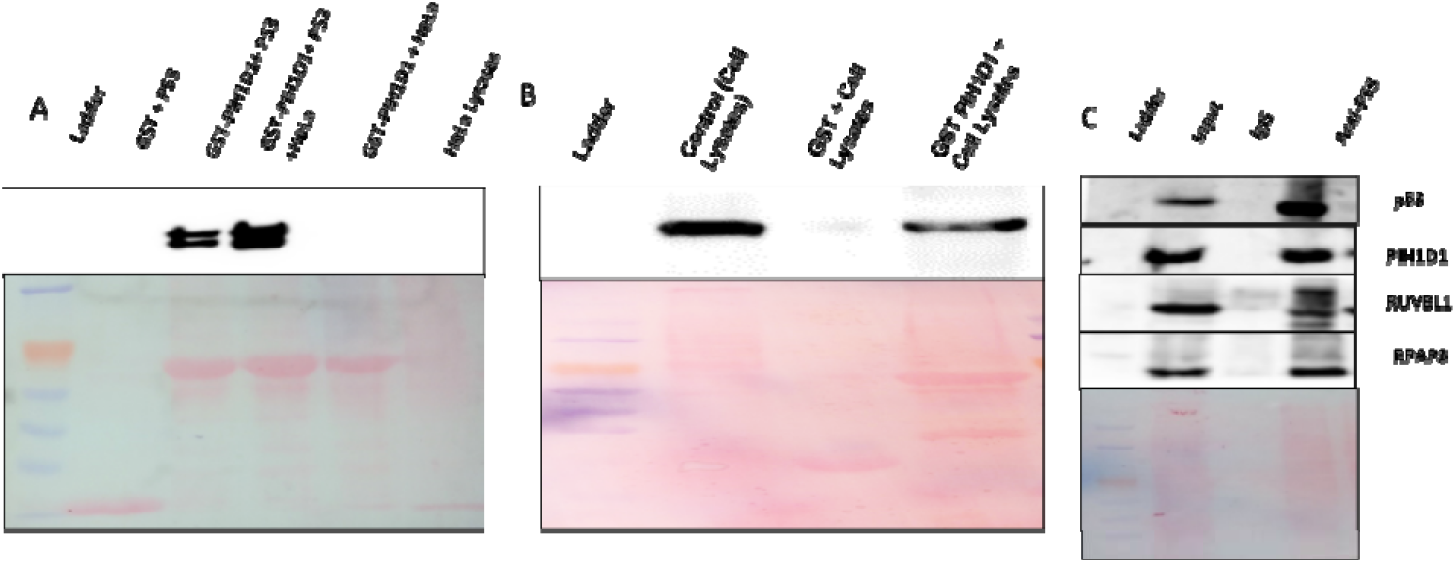
GST pull-down experiments showing PIH1D1 interacts p53 proteins physically in vitro. (A) GST pull-down experiments confirm the physical interactions between PIH1D1 and p53 proteins (B) GST tagged PIH1D1 was incubated with MCF-7 cell lysate the pulled down lysate was subjected to western blot with anti P53 antibody. Whole cell lysate and GST alone were used as control. (C) Co-IP with anti PIH1D1 antibody where 2mg/mL MCF7 cell lysates was incubated with 2ug antiPIH1D1 antibody.

Next to investigate the interaction of indigenous p53 with PIH1D1and other components of the R2TP complex, we incubated MCF 7 cell lysate with GST PIH1D1 as expected PIH1D1 was able to pull down indigenous p53 from MCF cell lysate (Figure 1B). Co-immunoprecipitation (Co-IP) assays were performed followed. The results demonstrated that P53 specifically interacts with PIH1D1 and so other components like RUVBL1, RUVBL2, and RPAP3, as these proteins were detected in the PIH1D1 immunoprecipitate (Figure 1C). Input lanes confirmed the presence of all proteins in total lysates, and Coomassie staining verified comparable protein loading. Collectively, these results confirm the successful expression and purification of functional recombinant PIH1D1 protein and validate the interaction of p53 with R2TP complex components.

### P53 interaction with PIH1D1 is independent of its C-Terminal domain

PIH1D1 is known to bind to DSDD/E motif on client proteins. P53 has a DSD instead of full DSDD motif on its C terminal domain (Figure 2 A). To assess the role of C-terminus of p53 in mediating the interaction with endogenous PIH1D1, we generated three GST tagged p53 constructs with and without C-terminus, and GST-tagged C-terminus of p53 only (Figure 2B), expressed and purified them (Figure 2C). The proteins were incubated with cell lysates of MCF7 and GST pull down assay was performed. Interestingly p53 with or without C terminal domain was able to interact with endogenous PIH1D1. However, the GST tagged C terminus of P53 was unable to pull down PIH1D1 from MCF7 cell lysates (Figure 2 D).

**Figure 2.**
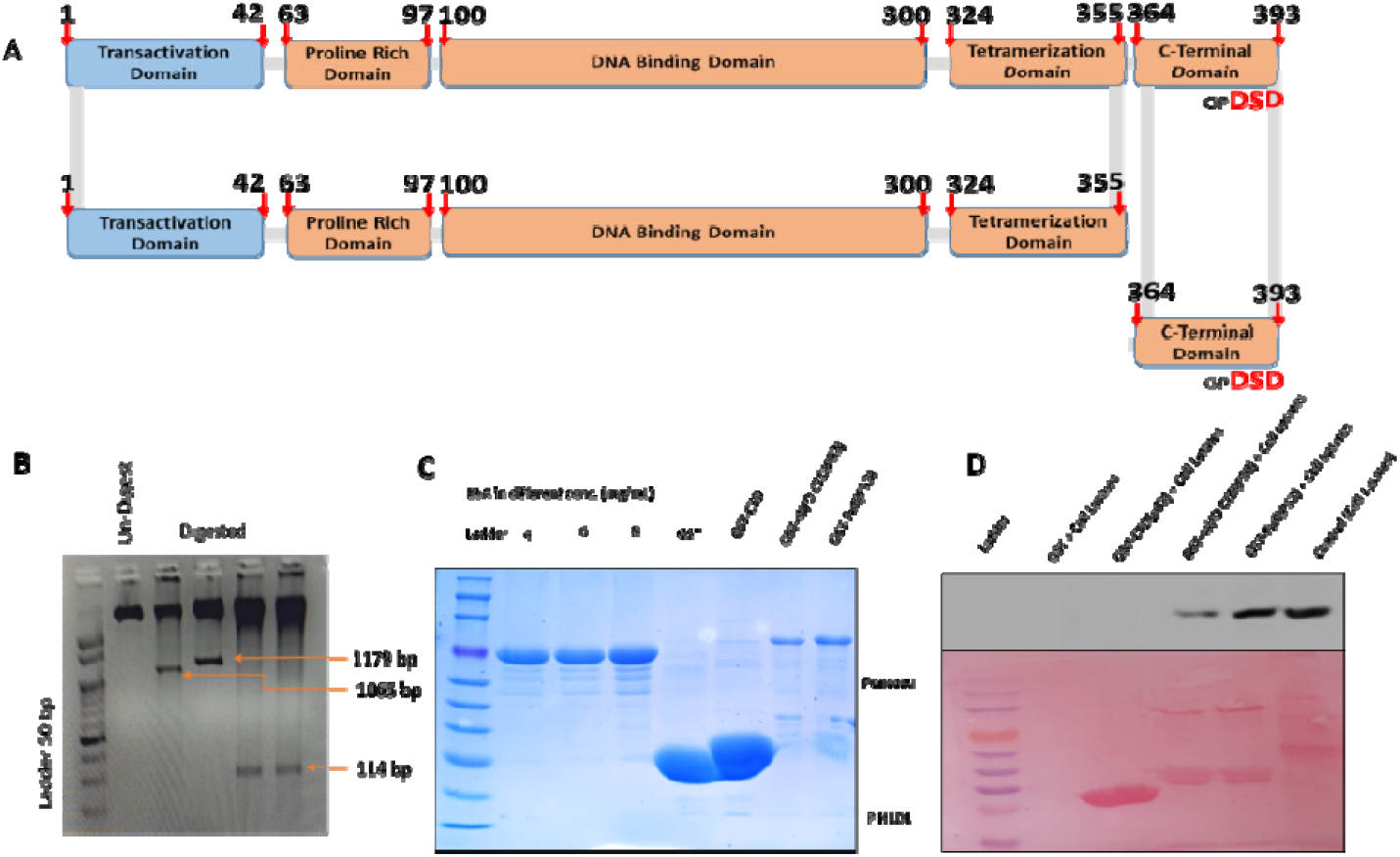
P53 C-terminus is dispensable for its interaction with PIH1D1. (A) Domain architecture of P53. (B) GST tagged p53 constructs with and without C-terminus, and GST-tagged C-terminus of p53, (C) Purified GST tagged c-terminus alone, gst-p53 without CTD and GST p53 with CTD (full length) quantified with respect to BSA and then 500 ng of purified was used for GST pull down (D)Interaction between endogenous PIH1D1 with GST p53, GST tagged C-terminal of p53 alone, truncated GST-tagged p53 without C-terminus and full length p53 was incubated with whole cell lysate of MCF7 cells. Whole cell lysate was used as control.

### PIH1D1 helps in the Stabilization of p53 protein

Taking into account, the established role of PIH1D1 as an adaptor protein within the R2TP co-chaperone complex, we sought to elucidate the mechanistic basis of its interaction with p53 specifically whether PIH1D1 contributes to p53 regulation. To assess the role of PIH1D1 in regulating p53 stability we try to knock out PIH1D1 with CRISPR Cas9 but due to very important role of PIH1D1 in the cells the Knockout cells were not survived. To address this issue we silenced PIH1D1 in MCF7 cells with two shRNA named shRNA 1 and ShRNA 2. Cells Lysate were collected from shRNA transfected cells and western blot was done for PIH1D1 and P53 respectively. PIH1D1 silencing led to substantial decrease in P53 protein levels in these cells (Figure 3A). We confirmed these results by silencing of PIH1D1 with siRNA (Figure 3B). Next,we investigated whether PIH1D1 silencing has any role in decreasing the half-life of p53, and we found PIH1D1 silenced cells have significant decrease in half-life of p53 as compared to control cells (Figure 3C). The density metric analysis (Figure 3D) was used to determine the half-life of p53.

**Figure 3.**
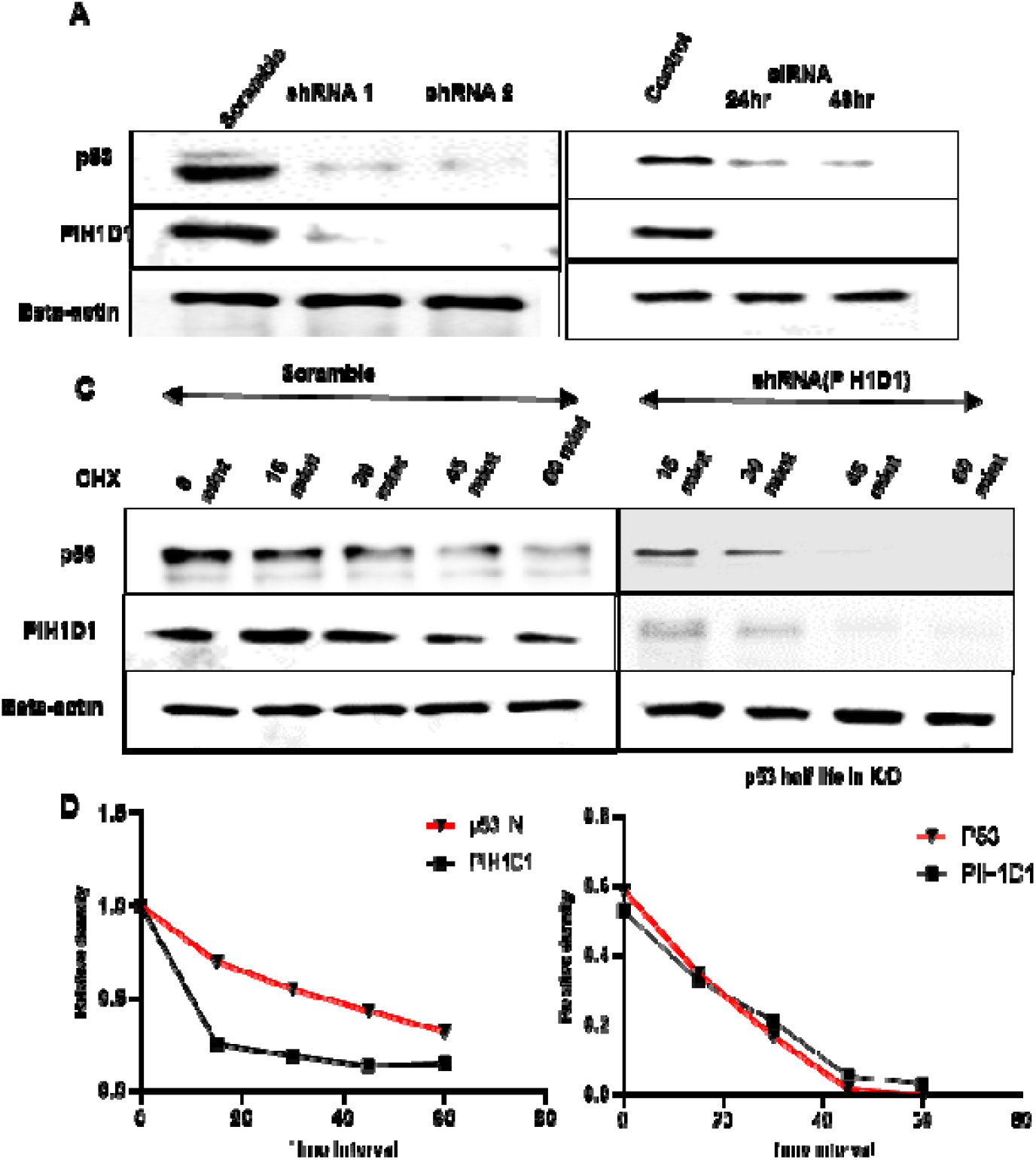
PIH1D1 is crucial for p53 stability. (A) MCF7 cells were transfected with two different shRNA 1 and 2, lysates were collected and immunoblotted with the indicated antibodies beta actin was used as internal control. (B) MCF-7 cells were transfected with siRNA and subjected to western blot with indicated antibodies. (C) PIH1D1 siRNA and scrambled siRNA transfected cells were treated with cyclohexamide and lysates were collected at different time intervals. These lysates were subjected to immunoblotting with indicated. (D) Half-life of P53 was calculated from the density metric analysis.

### PIH1D1 Silencing Reduces p21 but not p53 mRNA

In order to test does PIH1D1 silencing has any role in suppressing P53 transcription, we analyzed p53 mRNA in this cell line and we could not find any significant difference between mRNA levels in control vs PIH1D1 silenced cells (Figure 4A). We also quantified the p53 downstream target gene p21 levels in these cell as expected p21 expression was found significantly decreased in PIH1D1 silenced cells (Figure 4B). Comparative Gene expression analysis was shown significant for both gene p21 and p53. The expression of p21 and p53 were decrease in PIH1D1 knockdown cells as compare to control cells (Figure 4C). Data were normalize using GAPDH gene.

**Figure 4.**
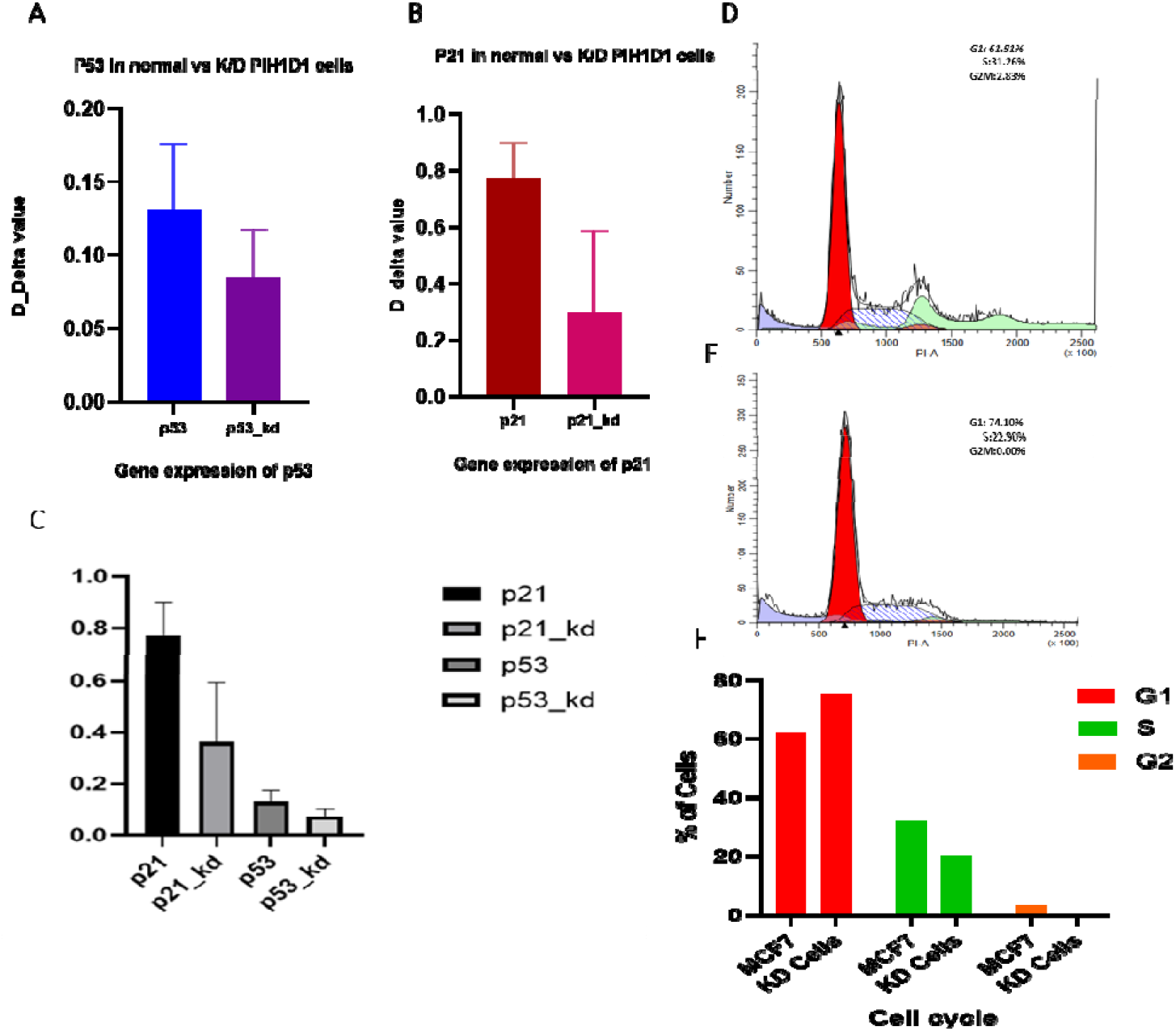
PIH1D1 knockdown alters p21 expression and Cell cycle. (A)Total cellular RNA isolated from PIH1D1 siRNA transfected cells was reverse transcribed and subjected to real time PCR for p53 and (B) p21 from same cells (18 S rRNA was used as an internal control). (C) Comparative Gene expression analysis for both gene p21 and p53. Cycle threshold (Ct) values were calculated for each PCR and relative fold change was calculated using 2DD Ct method. (D) Cell Cycle analysis by flow cytometry of control cells. (E) Cycle analysis of K/D (knockdown) for PIH1D1 cells. (F) The percentage of cells used for the plotting graph (G1 phase, middle S-phase and last phase represents G2/M phase).

### PIH1D1 Knockdown Impairs Cell-Cycle Progression via G1/S Phase Arrest

The outcomes of PIH1D1 knockdown (K/D) cells and control cells with normal PIH1D1 expression levels were contrasted. When PIH1D1 was silenced, flow cytometric profiles showed a clear change in cell-cycle progression.

In contrast to the control cells, PIH1D1 K/D cells showed a greater accumulation of cells in the G1 and S phases, suggesting a potential halt or delay in the G1/S transition (Figure 4E). The control cells, on the other hand, displayed a typical distribution throughout all cell-cycle phases and served as a baseline reference for normal PIH1D1 expression (Figure 4D). In order to show the relative alterations in cell-cycle dynamics between control and PIH1D1-silenced cells, the quantitative representation of the cell population in each phase (G1, S, and G2/M) was further displayed as a percentage graph (Figure 4F).

## Discussion

Upon successful completion of this study, we are now able to say that there is an entirely different novel mechanism for the interaction and stabilization of p53. Previously, the characteristic of p53 was found as p53 tetramerised [15–17]. One of the recently discovered HSP-90 co-chaperone complexes, named R2TP, also called PAQosome (particle for arrangement of quaternary structure), were speculated to play a role in the stabilisation of mutant p53[18,19]. The R2TP complex is considered the master regulator of cell growth and survival[12,20]. In our previous studies, we found that CKII-mediated phosphorylation occurs in the p53 protein at its specific site, DpSD [10,13]. While PIH1D1 is a more recently discovered gene with increased expression in multiple cancer types, p53 is also a well-known tumor suppressor that is commonly altered or dysregulated in various malignancies[21,22]. Comparative gene expression analysis showed in our earlier study that there was no transcriptional increase of this tumor suppressor gene, as evidenced by the fact that p53 transcript levels were unchanged in breast cancer (BRCA) tissues. However, compared to nearby normal controls, the PIH1D1 gene was significantly overexpressed in BRCA tissues, indicating a possible carcinogenic function of PIH1D1 that is not dependent on p53 transcriptional regulation [12]. Furthermore, we observed that the p53 and PIH1D1 genes were overexpressed in GBM tissues as compared to normal tissues, similar expression was found in the recent studies [23]. In our present study the GST pull-down was done for p53 protein, we observed that GST-PIH1D1 was able to pull down pure p53 both alone as well as from the cell lysate of Hela cells in which purified p53 was incubated (Figure 1A). Furthermore we investigated the interaction of indigenous p53 with PIH1D1 and other components of the R2TP complex, and we observed that PIH1D1 was able to pull down indigenous p53 from MCF cell lysate (Figure 1B). Co-immunoprecipitation (Co-IP) assays results demonstrated that P53 specifically interacts with PIH1D1 and so other components like RUVBL1, RUVBL2, and RPAP3, as these proteins were detected in the PIH1D1 immunoprecipitate (Figure 1C). Collectively, results confirmed the expression and purification of functional recombinant PIH1D1 protein and validate the interaction of p53 with R2TP complex components. PIH1D1 is known to bind to DSDD/E motif on client proteins[24]. Previous study has revealed P53 has a DSD instead of full DSDD motif on its C terminal domain[25–27]. To check the role of C-terminus of p53 in mediating the interaction with endogenous PIH1D1 (Figure 2 A), we generated three GST tagged p53 constructs with and without C-terminus, (Figure 2B), and purified them (Figure 2C). Interestingly in our study, we found p53 with or without C terminal domain interact with endogenous PIH1D1. However, the GST tagged C terminus of P53 was unable to pull down PIH1D1 from MCF7 cell lysates (Figure 2 D).

Furthermore to established role of PIH1D1 as an adaptor protein within the R2TP complex, we elucidated the mechanistic basis of its interaction of PIH1D1 with p53 in the p53 regulation. We silenced PIH1D1 in MCF7 cells with two shRNA named shRNA 1 and ShRNA 2, and we observed that PIH1D1 silencing led to substantial decrease in P53 protein levels in these cells (Figure 3A). We confirmed these results by silencing of PIH1D1 with siRNA (Figure 3B). In the next step, we investigated the half-life of p53 in normal cells and knockdown cells and found PIH1D1 silenced cells have significant decrease in half-life of p53 (Figure 3C) as compared to control cells. The half-life of P53 was calculated from the density metric analysis (Figure 3D). Since there were no discernible differences in the expression of p53 mRNA between control and PIH1D1-silenced cells, these findings collectively suggest that PIH1D1 does not affect p53 at the transcriptional level. Instead of directly controlling transcription, the R2TP complex mainly regulates client proteins post-translationally[5,28]. The R2TP complex, composed of PIH1D1, RPAP3, and the AAA+ ATPases RUVBL1 and RUVBL2, acts as a co-chaperone for HSP90, facilitating the assembly and stabilization of large macromolecular complexes [29]. Through phosphorylation-dependent interactions and chaperone-mediated stabilization, prior research has shown that R2TP and its associated proteins aid in the maturation and stability of a number of regulatory proteins, including PIKK family kinases (like mTOR and ATM) and elements of the RNA polymerase II machinery[30,31]. The substantial reduction in p21 (CDKN1A) transcript levels, a known downstream target of p53 that triggers cell-cycle arrest at the G1/S checkpoint, suggests that PIH1D1 silencing affects p53 functional activity rather than transcription [32]. Prior research has shown that certain post-translational changes, such as phosphorylation, acetylation, and ubiquitination, tightly regulate p53 function and impact its stability, nuclear localization, and DNA-binding ability[33,34]. PIH1D1 silencing appears to impair p53 functional activity rather than transcription, as evidenced by the significant decrease in p21 transcript levels, a known downstream target of p53. This suggests that the stability or activation potential of p53 may be impacted by post-transcriptional or post-translational regulation by PIH1D1. In line with this finding, PIH1D1 knockdown (K/D) cells’ cell-cycle distribution was significantly altered by flow cytometric analysis when compared to control cells that expressed normal amounts of PIH1D1. A cell-cycle arrest or delay at the G1/S transition was specifically shown by the higher accumulation in the G1 and S phases of PIH1D1-depleted cells[35–37]. The normal phase distribution of control cells, however, is a sign of typical proliferative behavior. Further evidence for PIH1D1’s role in maintaining proper cell-cycle progression through p53–p21 signaling comes from quantitative analysis of cell populations during the G1, S, and G2/M phases (Figure 4F). All things considered, our results lend credence to the idea that PIH1D1 positively controls p53 stability and downstream transcriptional activity, and that its deficiency results in decreased p21 expression, dysregulated cell-cycle progression, and impaired p53 signaling.

Upon successful completion of this study, we are now able to conclude that an entirely different novel mechanism has been found for the interaction and stabilization of p53. Upon interaction between p53 and PIH1D1 protein, p53 is stabilized by PIH1D1 protein, a R2TP complex member, and this interaction was independent of the C-terminal domain of p53 protein. We have also observed that p53 protein levels were affected after the alteration in expression levels of PIH1D1, a component of the R2TP complex, as confirmed in vivo using MCF7 cell lines. On the basis of the finding, we suggest that P53 protein is stabilized and remodelled by the co-chaperone protein R2TP complex. Presumably, this new mechanism will serve as a flashpoint to restore/activate the function of p53 in different cancer cells, in which it leads the control of the cause of cancer and cell cycle progression.

## Acknowledgement

Authors wish to thank all Lab members of Dr. Riyaz’s Lab, Dept of Biochemistry, AIIMS, and New Delhi for his technical support and valuable contributions to this project. We also wish to acknowledge all the authors for preparing and editing the manuscript and funding agencies (CSIR and ICMR).

## Authors contributions

R.A.M conceived the project. D.K. implemented and tested the programs. Q.A, R.A.M and D.K read, edited and prepared the manuscript. All authors approved the final manuscript.

## Consent statement

Consent statement is not applicable

## Data availability

The data that support the findings of this study are openly available in bioRxiv with reference number BIORXIV/2026/704515 and the repository URL as https://www.biorxiv.org/content/10.64898/2026.02.07.704515v1.

## Ethics approval and consent to participate

Ethics was taken from Institutional committee on biosafety for recombinant DNA research, All India institute of Medical Sciences, New Delhi. (REF.No. IBSC 1221_DK_RM)

